# c-Kit mediates cutaneous sensory axon innervation and multi-kinase inhibitor-induced neurotoxicity

**DOI:** 10.1101/2021.09.13.460106

**Authors:** Adam M. Tuttle, Matthew B. Pomaville, Katherine C. Delgado, Kevin M. Wright, Alex V. Nechiporuk

## Abstract

Peripheral somatosensory neurons innervate the skin and sense the environment. Loss of skin innervation, often caused by the “dying back” of distal somatosensory axons, is a common side effect of drug-induced peripheral neuropathies (DIPNs) and results in pain and sensory dysfunction. Targeted cancer therapies frequently employ multi-kinase inhibitor (MKI) drugs that each block multiple receptor tyrosine kinases. Many MKIs produce DIPNs but the molecular targets and cellular mechanisms underlying these are unknown. We performed live-imaging of cutaneous somatosensory axons in larval zebrafish during treatment with several MKIs known to induce DIPNs, and observed axonal retraction consistent with a dying back pathology. These results were replicated in mouse somatosensory neurons. Genetic knockout of potential MKI targets identified c-Kit receptor as a regulator of sensory axon innervation and a major target of these MKIs mediating loss of axonal density. In both fish and mammals, Kit receptor is expressed in cutaneous somatosensory neurons and its ligand, Kitlg, is expressed in the skin. Mosaic misexpression of Kitlg in the skin induced dramatic increases in local sensory axon density, suggesting an important role for Kit signaling in cutaneous axon growth and maintenance. Immunostaining and structure-function analysis revealed Src, a downstream Kit target, mediates Kit’s role in cutaneous axon innervation and MKI neurotoxicity. Our data shows that the Kit-Src signaling pathway has a major role in cutaneous sensory axon innervation and is a potential therapeutic target to address DIPNs caused by MKIs and other compounds.

## INTRODUCTION

Personalized cancer drug therapies centered around multi-kinase inhibitor (MKI) drugs are an effective and increasingly common form of cancer treatment. Each MKI targets multiple kinases that regulate cancer cell proliferation and growth. MKIs are a rapidly expanding class of drugs, however, many MKIs can cause damage to the peripheral nervous system, termed drug-induced peripheral neuropathies (DIPNs)^1-4^. DIPNs can cause sensory dysfunction and pain that significantly impacts patient quality of life and can limit dosage and duration of otherwise effective drug therapies, thereby negatively impacting patient outcomes^5^. Many MKIs induce peripheral neuropathy in >10% of patients as well as a host of likely-related adverse effects such as musculoskeletal pain, headaches, and plantar-palmar pain^2-4,6,7^. To date, there are no specific therapies to address DIPNs, whether caused by MKIs or other compounds, and they remain a major clinical complication of cancer drug treatments.

The molecular and cellular causes of DIPNs from “traditional” chemotherapy drugs, such as vinca alkaloids and taxanes are under intense investigation. The cellular etiology of these DIPNs is the progressive dying-back of distal axon terminals of small-fiber, unmyelinated sensory neurons in the extremities^8,9^. This results in a reduction of cutaneous axon innervation in these areas and subsequent pain and numbness. The molecular cause of this axon die-back varies based on compound and ranges from changes in microtubule dynamics (vincristine, taxanes) to disruption of proteolysis (bortezomib). Whether this stereotypical axon die-back and changes in cutaneous axon density are the cause of MKI-induced peripheral neuropathies is unknown. The predominant target of MKIs are receptor tyrosine kinases (RTKs). Thus, it is likely that the neurotoxicity of many MKIs is mediated through the inhibition of specific RTKs or RTK families. But which RTKs mediate neurotoxicity and how each RTK may specifically affect neuronal health and cellular behavior remains to be elucidated.

Each MKI inhibits multiple RTKs at various affinities, creating a unique inhibition profile. One such RTK is c-Kit, a transmembrane Type III RTK. It is activated upon binding of its primary ligand, Stem Cell Factor/Kit-Ligand (SCF/KITLG). C-Kit receptor is expressed in a subset of mammalian dorsal root ganglia (DRG) neurons in embryos and adults and SCF is expressed in rodent adult skin^10-12^. Several studies of embryonic and adult mammalian systems have suggested possible roles for c-Kit signaling in DRG neurons^11,13-15^, though its specific function is still unclear and a role regulating cutaneous innervation of somatosensory neurons has not been established.

One downstream target of c-Kit with roles in axon outgrowth and regeneration is the Src Family Kinases (SFKs). SFKs are a group of non-receptor tyrosine kinases that regulate signal transduction of many cell membrane receptors. Src is a single member of the family; it binds to activated c-Kit receptor^16,17^ to mediate its downstream signaling. Activated Src/SFKs mediate axon outgrowth and guidance in *Xenopus*, chick, and *C. elegans*^*18-20*^. Src/SFKs have been shown to promote axon regeneration in motor neurons in *C. elegans*^*21*^, sciatic nerve in rats^22^, and zebrafish somatosensory neurons^23^. Much of this role is likely due to local Src activity in the tips of axon growth cones, where it regulates local tyrosine kinases that, in turn, impact local filopodial/adhesion dynamics^18^. However, the link between Kit signaling and SFK activity in peripheral axons has not been explored, and how Src activity may relate to MKI DIPNs is unknown.

In this study, we have developed a novel zebrafish model to examine the effect of several MKIs that cause DIPNs in human patients (sorafenib, ponatinib, and dasatinib). We use zebrafish larvae as they are amenable to a combination of high-throughput drug delivery, exquisite imaging access, and genetic interrogation. Zebrafish larvae also have functional somatosensory neurons that innervate the skin that are homologous to those affected in DIPNs^24-27^. We find that treatment with MKIs or the neurotoxic chemotherapy drug vincristine induces loss of distal cutaneous axon density and retraction of axon terminals. This effect is consistent with the stereotypical axonal “die-back” pathophysiology of DIPN progression. We find similar neurotoxic effects in cultured mouse DRG neurons. We identify c-Kit receptor (*kitb* in zebrafish) as an MKI target mediating the observed neurotoxicity. Genetic loss of *kitb* leads to a similar loss of cutaneous axon density, while hyperactivation of Kitb signaling induces dramatic, highly-localized increases in axon density. We find that Src/SFK activity downstream of Kitb is required for maintaining normal axonal density. Treatment with MKIs induced significant loss of SFK activity in distal axons while local hyperactivation of Kitb signaling increased axonal SFK activity. Thus, we establish c-Kit-Src signaling as an important promoter of cutaneous axonal innervation and maintenance, as well as a mediator of the neurotoxicity of certain MKIs. These findings highlight the power of our *in vivo* technical approach to address the neurobiology underlying an underserved clinical complication of MKI drugs; it also opens up new avenues of research into novel cellular roles for signaling pathways in axonal development and disease.

## RESULTS

### MKI treatment induces loss of cutaneous sensory axon innervation

We first asked if MKIs affect cutaneous somatosensory axon density in zebrafish. Zebrafish larvae possess Rohon-Beard neurons (RBs), unmyelinated somatosensory neurons that innervate the larval skin. These neurons are homologous in function and expression profile to the small-fiber sensory neurons of the dorsal root ganglia (DRG) affected in many human DIPNs^24-27^. By 48 hours post-fertilization (hpf), RB sensory neurons are functional and their axon arbors fully innervate the larval skin and maintain this functionality for weeks^28,29^. Zebrafish develop DRG neurons that will replace RBs as the primary somatosensory neurons innervating the skin, however, this transition takes place much later in development during the formation of bony scales (1-2 months post-fertilization)^29^. To visualize cutaneous RB axons *in vivo* we used larvae carrying both the *SAIGFF213A* (a GAL4 driver that labels distinct types of neurons, including the RBs^30^) and a UAS:GFP reporter transgenes (this double transgenic line hereafter referred to as *RB:GFP*). By 48 hpf, GFP+ RB axons have reached the tail tip, arborized, and extended along the perimeter of the caudal tail edge. In order to evaluate the potential neurotoxic effect of MKIs on established distal axon terminals and arbors, we chose 72 hpf as the beginning of the drug treatment window after cutaneous distal axon outgrowth and arborization had stabilized.

We treated *RB:GFP* larvae with several MKIs that induce DIPNs in human patients (sorafenib, ponatinib, dasatinib) at 72 hpf for 48 h, then imaged the GFP-labeled axons of the distal caudal tail tip (Fig. 1A,B). As a positive control, we also treated with vincristine, a commonly used vinca alkaloid chemotherapy drug that induces peripheral neuropathy in patients, largely due to the “dying back” of distal axon terminals. To quantify axonal density within the thin skin of the distal tail, we used ImageJ to create a maximum intensity Z-stack projection and measured the proportional area of a 50×100 μm region of interest at the tip of the tail occupied by GFP-labeled axons. Treatment with vincristine induced loss of distal cutaneous axon density (Fig. 1C,G), a phenotype that is consistent with results in human patients^8^ and mammalian models^31^. Using this assay, we next found that MKIs induced significant, dose-dependent loss of distal cutaneous axon density (Fig. 1D-F,H-J, Suppl. Fig. 1A-C) when compared to vehicle treatment. The reduction of distal axon density induced by vincristine and these MKIs indicates a substantial degree of sensory neurotoxicity. However, unlike vincristine, the cellular bases of MKI-induced human peripheral neurotoxicity are not known.

**Figure 1:**
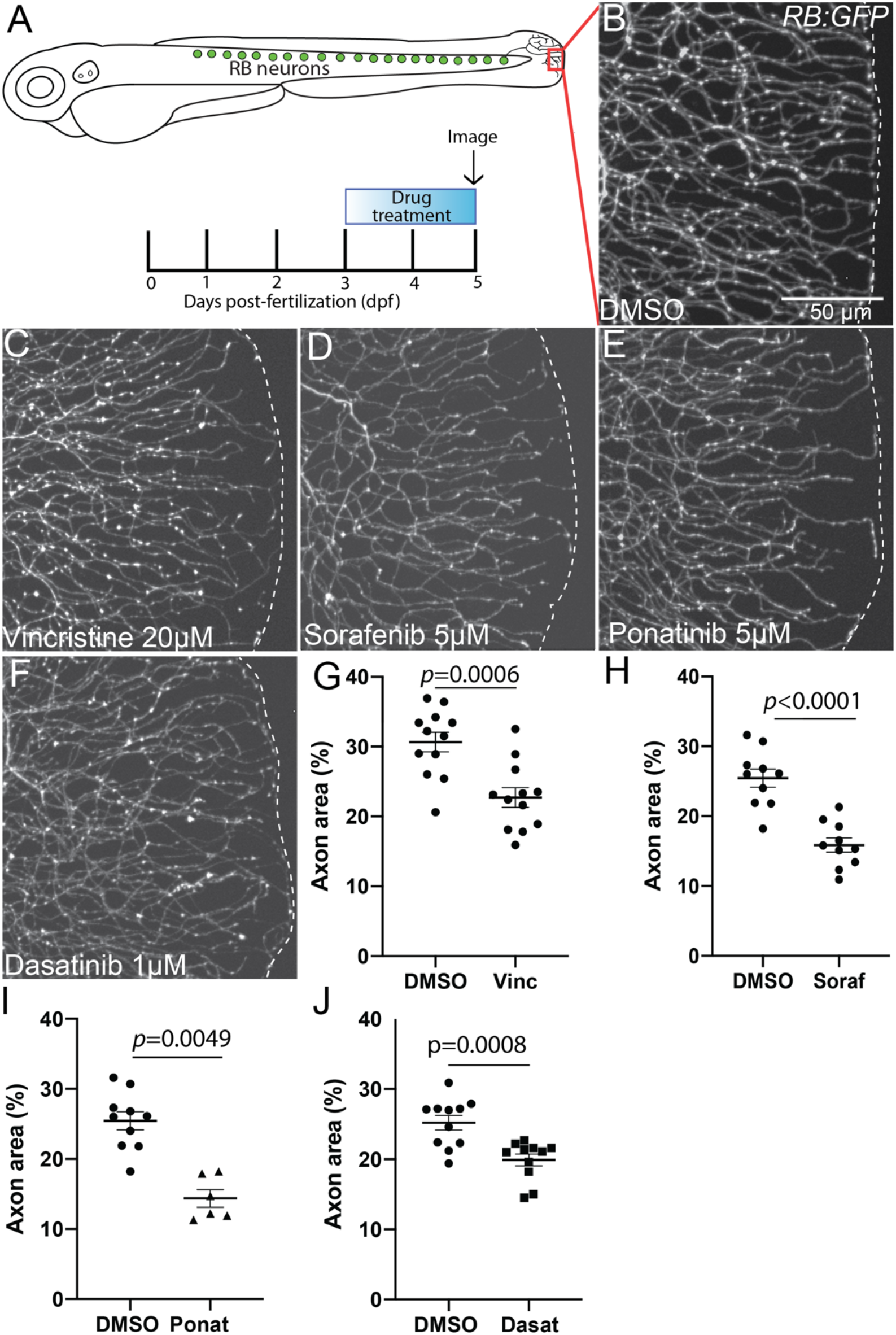
Treatment with multi-kinase inhibitors or the neurotoxic chemotherapy drug vincristine induce loss of cutaneous axon density in zebrafish caudal tail. (A) Diagram of Rohon-Beard neurons, drug treatment and imaging paradigm, and imaging location (red box). (B-F) Live images of somatosensory axons at the caudal tail tip labeled by *RB:GFP* after 48 h treatment. Several MKIs produced significant loss of cutaneous axon density, as well as chemotherapeutic drugs with known peripheral neurotoxicity (white dashes indicate tail edge). (G-J) Quantification of loss of distal axon density, analyzed by unpaired T-test. (G) DMSO=30.7±1.4 % area vs. vincristine=22.7±1.4. (H) DMSO=25.4±1.3 % area vs. sorafenib=15.9±1.0. (I).DMSO=28.3±3.3 % area vs. ponatinib=17.3±1.3. (J) DMSO=25.2±1.1 % area vs. dasatinib=19.9±0.9. Error bars represent SEM.

### Axon terminal retraction underlies MKI-induced axon loss

To determine the cellular basis for the loss of cutaneous axon density, we performed time-lapse imaging of sparsely-labeled RB axons in the caudal tail. Injection of a UAS:mCherry plasmid at one-cell stage in *RB:GFP* embryos mosaically labeled RB neurons and allowed imaging of axon terminals in the caudal tail from individual neurons during drug treatment. Control RB axon terminals are dynamic and often extend, retract, and explore over time (Movie 1, Fig. 2A). Compared to vehicle treatment, application of vincristine, sorafenib, and ponatinib induced notable retraction of individual axon terminals over a period of 10 h, though terminal behavior was still dynamic (Movies 2-4, Fig 2B-D). To analyze the movement of these time-lapse imaged individual axon terminals, we performed kymograph analysis on distal terminals. Kymographs show consistent proximal movement of terminals during drug treatment (Fig. 2E-H). Quantification of the net movement of distal axon terminals over time found that control terminals exhibited minor net terminal extension over 10 h, while sorafenib, ponatinib, and vincristine treatment all induced significant net retraction from the initial position (DMSO=1.5±1.1 μm net distal movement, sorafenib=-6.6±0.7, ponatinib=-4.4±1.2, vincristine=-6.7±1.4). The net movement of drug-treated axon terminals proximally, as opposed to remaining stationary, suggests significant retraction as opposed to a failure of outgrowth. It is also possible that distal axonal arborization or growth between 3 and 5 dpf was affected by the MKI treatments. However, repeated measures of distal axon density of individual control larvae during the drug treatment window indicated no significant changes in distal tail axon density from 3 to 5 dpf (Fig. 2J) or 4 to 5 dpf (Fig. 2K). Taken together, these data indicate that the loss of axon density following MKI treatment is due to axon terminal retraction/die-back and are consistent with the axon die-back found in many other DIPNs.

**Figure 2:**
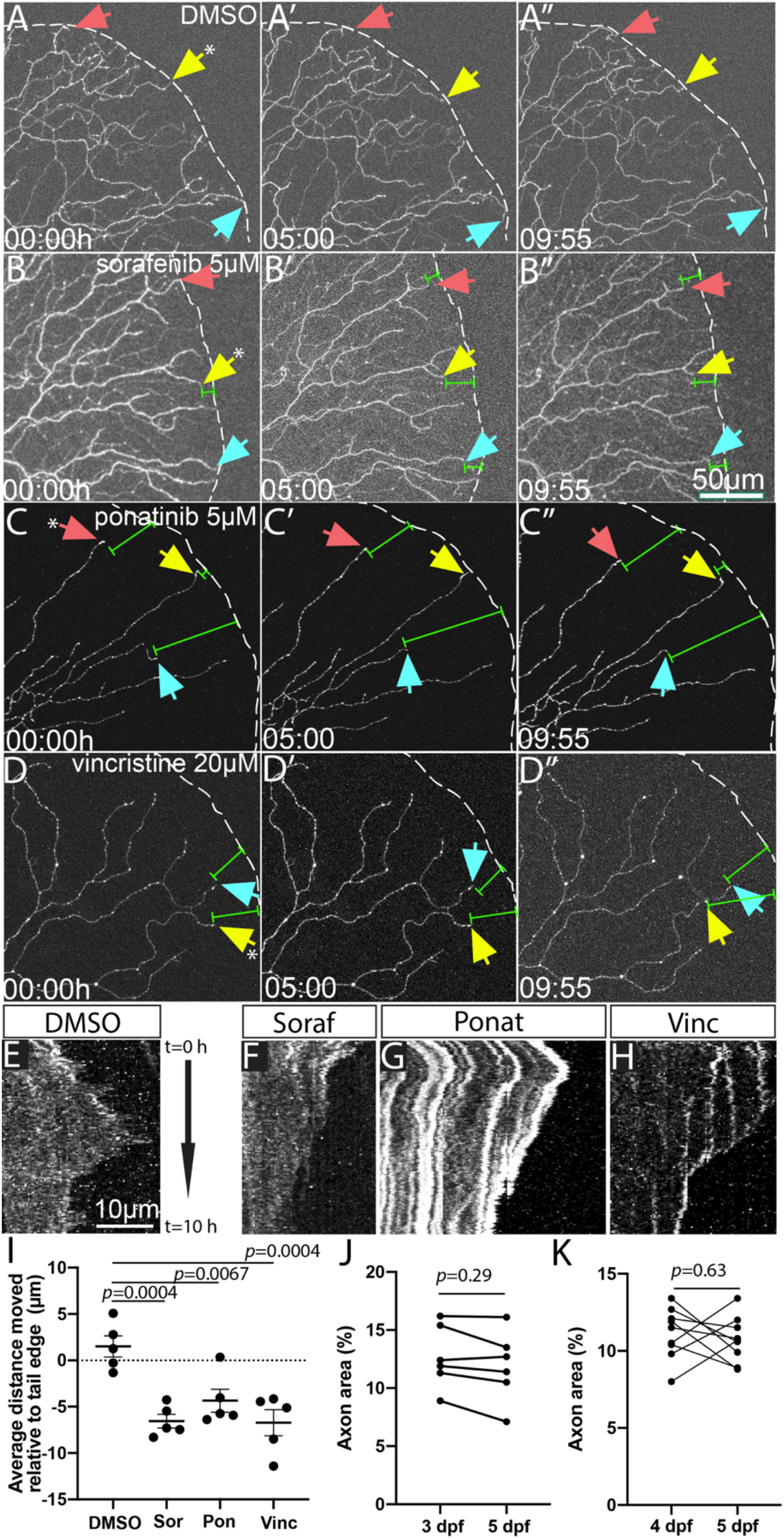
Time lapse imaging reveals axon terminals retraction following sorafenib, ponatinib, and vincristine treatment. (A-D) Stills from Movies 1-3, respectively, of distal axons of sparsely labeled Rohon-Beard neurons. Over 10 h, sorafenib, ponatinib and vincristine treatment induced retraction and degeneration events of distal axon tips. (white dashes = tail edge, arrows indicate specific axon terminals followed over time, green measurement bars indicate distance from edge. (E-H) Kymographs of individual axon terminals (indicated by asterisk in panels A-D) from respective movies. DMSO terminals (E) show dynamic extension and retraction, but sorafenib, ponatinib, or vincristine treatment induce notable net retraction of axon terminals. (I) Quantification of axon retraction in 10 h time lapse movies. Compared to DMSO-treated larvae that had a net increase in axon outgrowth, sorafenib, ponatinib, and vincristine treatment induced a significant net retraction of distal RB axon terminals in labeled neurons by one-way ANOVA with post hoc Dunnett’s test. DMSO=1.5±1.1 μm, vs. sorafenib=-6.5±0.7, ponatinib=-4.4±1.2, vincristine=-6.7±1.4. Error bars represent SEM. (J,K) Repeated measures of axonal density of wildtype control larval tails analyzed by paired T-test from 3-5 dpf (J) or 4-5dpf (K). (J): 3 dpf=12.3±1.1%, 5 dpf=11.9±1.2. (K) 4dpf=11.1±0.6%, 5dpf=10.7±0.5.

### Loss of Kitb receptor phenocopies drug treatment

The mechanisms by which MKIs induce neurotoxicity at a molecular level are not known. A possible cause of neurotoxicity is inhibition of one or more kinases targeted by these MKIs that ultimately leads to axon die-back and loss of axon density. MKIs can target a range of kinases, but RTKs are often the primary targets of many MKIs. We searched the literature for the common RTK targets of sorafenib, ponatinib, and dasatinib that are reported to be highly inhibited (IC^50^≤100 nM *in vitro*)^32-36^. Among these RTKs, c-Kit and Ret are specifically expressed in zebrafish RBs^37,38^, and c-Kit is a major inhibitory target of all three MKIs. Furthermore, c-KIT and RET are reported to be expressed in mammalian nociceptive neurons with roles in neurodevelopment, regeneration, and/or pain sensation^10,13,39,40^. We chose these receptors as candidates for RTKs that may underlie neurotoxicity of these MKIs and examined them for roles in zebrafish RB neurons.

Zebrafish have two paralogues of c-Kit, Kita and Kitb, but only Kitb is specifically expressed in the RB neurons^41^. In *RB:GFP* larvae, transient CRISPR-Cas9 mediated knockdown of Kitb induced a significant loss of caudal tail axon density at 3 and 5 dpf (Fig. 3A). Notably, knockdown of the other candidate expressed in RBs, Ret, had no effect on somatosensory axons in the caudal tail (Fig. 3A). To confirm the transient Kitb knockdown phenotype, we obtained a zebrafish mutant line (*kitb*^*sa15348*^), containing a premature stop in *kitb* (Y197X). *RB:GFP* transgenic larvae homozygous for the *kitb* mutant allele had significantly reduced tail cutaneous axon density at 5 dpf compared to wild-type siblings (Fig. 3B-D, WT = 13.4±0.6 vs. *kitb* mutant=7.9±0.7 %axonal area). Caspase-3 immunolabeling and live imaging analysis indicated that the reduction in cutaneous axon density was not due to premature RB neuron death (Suppl. Fig. 2). Injection of 25 pg of *kitb-p2a-mCherry* mRNA into one-cell stage *kitb* mutant or sibling embryos significantly suppressed the loss of tail innervation in *kitb* mutants by 3 dpf (Fig. 3D), demonstrating that the loss of *kitb* is indeed responsible for this phenotype. These data indicate Kitb is a regulator of cutaneous somatosensory axon innervation in zebrafish larvae.

**Figure 3:**
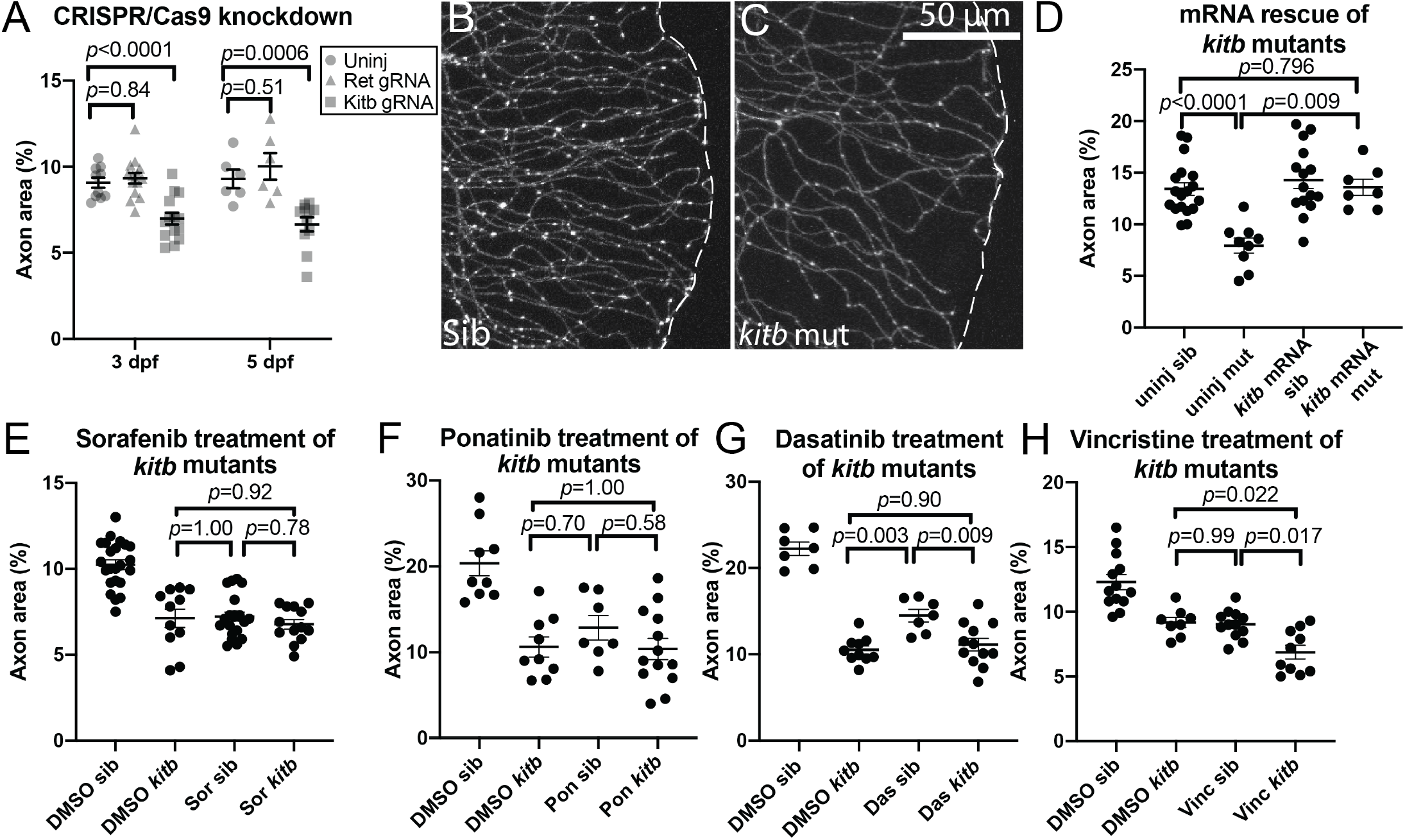
*kitb* is a major target mediating neurotoxicty of several MKIs. (A) Quantification of distal cutaneous axon density in caudal tail after CRISPR-Cas9-mediated knockdown of either *ret* or *kitb*. 3 dpf: uninjected=9.1±0.3% area, *ret* gRNA= 9.3±0.3, *kitb* gRNA=7.0±0.3. 5 dpf: 9.3±0.5%, *ret* gRNA= 10.3±0.9, *kitb* gRNA=6.7±0.4. (B,C) Live imaging at 5 dpf of *kitb* homozygous mutants or siblings (sib) in the *RB:GFP* transgenic background (white dashes = tail edge). (D) Expression of *kitb* mRNA suppresses *kitb* mutant axon density defect. Uninjected sib=13.4±0.6 % area, uninjected mutant= 7.9±7, *kitb* mRNA sib=14.9±0.9, *kitb* mRNA mutant= 13.6±0.8. (E-G) Quantification of *kitb* mutants treated with either sorafenib (E), ponatinib (F), or dasatinib (G) using the same treatment paradigm from Fig. 1A. While treatment with either drug or genetic loss of *kitb* induced loss of cutaneous axon density, treatment of *kitb* mutants with drug did not exacerbate this phenotype, indicating Kitb is a major target underlying their peripheral neurotoxicity in this case. (E) DMSO sib=10.2±0.3 % area vs. DMSO *kitb*=7.1±0.5, *p*<0.0001; vs. sorafenib sib=7.2±0.3, *p*<0.0001; vs. sorafenib *kitb*=6.8±0.3, *p*<0.0001. (F) DMSO sib=20.4±1.5 % area vs. DMSO *kitb*=10.6±1.2, *p*<0.0001; vs. ponatinib sib=12.9±1.4, *p*=0.0051; vs. ponatinib *kitb*=10.4±1.2, *p*<0.0001. (G) DMSO WT=22.2±0.8% vs DMSO *kitb*=10.5±0.5, *p*<0.0001; vs. dasatinib WT=14.5±0.7, *p*<0.0001; vs. dasatinib *kitb*=11.1±0.7, *p*<0.0001. (H) Quantification of vincristine treatment of *kitb* mutants. In contrast to sorafenib and ponatinib, vincristine treatment significantly exacerbates axon density loss in *kitb* mutants. DMSO sib=13.3±0.6 %area vs. DMSO *kitb* mutant=9.2±0.4, *p*=0.0006; vs. vincristine sib=9.0±0.3, *p*<0.0001; vs. vincristine *kitb*=6.9±0.5, *p*<0.0001. Error bars represent SEM. Two-way ANOVA with post hoc Tukey’s HSD test for data sets.

### Kitb is a major neural target of MKIs

In order to test if Kitb is a major target that mediates peripheral neurotoxicy, we treated *RB:GFP kitb* mutant or sibling larvae with either vehicle, sorafenib, ponatinib, or dasatinib. If Kitb primarily mediates the observed sorafenib-, ponatinib-, or dasatinib-induced axonal loss, we expected genetic removal of Kitb would preclude additional neuropathic effects from drug treatment. In accordance with this, we observed no significant exacerbation of the axon density loss at 5 dpf in *kitb* mutants treated with any of the individual MKIs compared to vehicle-treated *kitb* mutants (Fig. 3E-G). There was also no significant difference between vehicle-treated *kitb* mutants and sorafenib/ponatinib-treated wildtype larvae, though dasatinib-treated wildtype larvae had less axon density reduction than vehicle- or dasatinib-treated *kitb* mutants. In contrast, treatment of *kitb* mutants with vincristine significantly increased the axon density loss (Fig. 3H). Altogether, these data suggest that Kitb is a specific, major target of sorafenib, ponatinib, and dasatinib that mediates their peripheral neurotoxicity. Furthermore, loss of Kit signaling may exacerbate the neurotoxicity of other chemotherapeutic compounds.

We next asked if these three drugs had a similar effect on Kit-positive neurons in mammalian DRGs. Dissected embryonic mouse DRG explants cultured in media containing high levels of the Kit-ligand SCF (but without other exogenously supplemented growth factors such as NGF) survive and produce a high proportion of c-Kit+ neurons^12,42^. DRGs were dissected (E15.5) and cultured for 24 h in media containing SCF ligand, then treated for 24 h with MKIs. Treatment of cultures with sorafenib, ponatinib, and dasatinib led to significantly shorter neurites and reduced axonal density compared to vehicle (Fig. 4A-H). These data suggest MKIs have a similar effect on mammalian somatosensory neurons as we observe in zebrafish.

**Figure 4:**
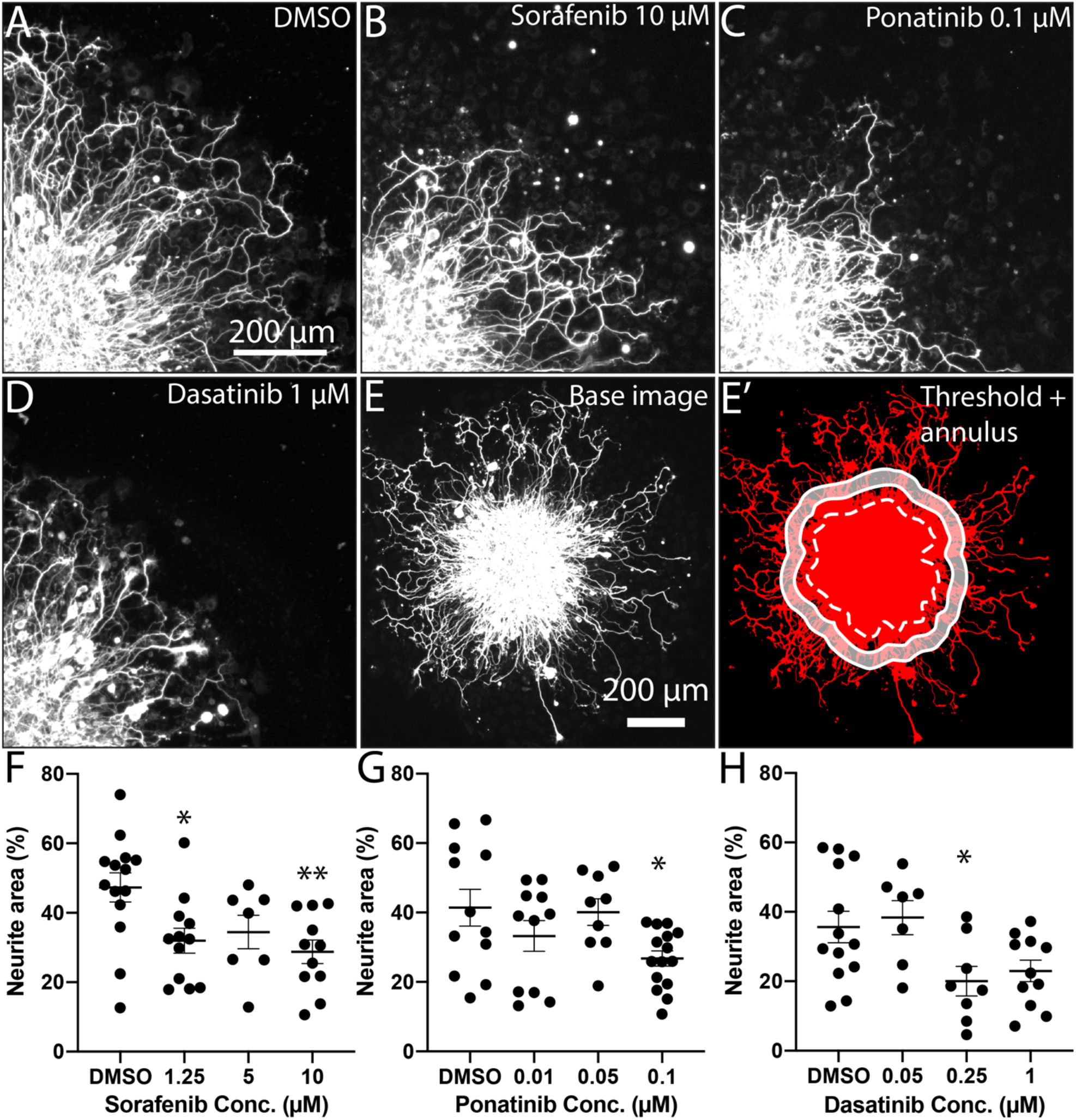
MKI treatment induces loss of neurite density in mammalian DRGs. (A-D) Single quadrant images of fixed DRG explants immunostained for βIII-Tubulin treated with vehicle or MKI compounds. MKI treatment led to shorter and less dense neurites. (E,E’) Demonstration of neurite area analysis. Immunostained DRG explants were imaged (E), thresholded (E’, red), and then the explant core was identified (dotted white line) and a 50 μm thick annulus (white shaded area) was created by drawing the inner and outer perimeter 50 and 100 μm from the edge of the explant core, respectively. (F-G) Quantification of neurite area, analyzed by one-way ANOVA with post hoc Dunnett’s test. (F) Sorafenib: DMSO=47.3±4.2% area, 1.25 μM 32.0±3.6, *p*=0.016, 5 μM 34.5±4.8 *p*=.116, 10 μM 28.8±3.4, *p*=0.0039. (G) Ponatinib: DMSO=41.4±5.3, 0.01 μM 33.3±4.4, *p*=0.353, 0.05 μM 40.1±3.9, *p*=0.99, 1 μM 26.8±2.2 *p*=0.0215 (H) Dasatinib: DMSO=35.6±4.5, 0.05 μM 38.4±4.9, *p*=0.954, 0.25 μM 20.0±4.3, *p*=0.038, 1 μM 23.0±3.1, *p*=0.073. Error bars represent SEM. ^*^=*p*<0.05, ^**^=*p*<0.01.

### Kit ligand b promotes local cutaneous axon innervation

*c-Kit* is expressed in a subset of mammalian DRG neurons, largely peptidergic unmyelinated C-fibers^10^ and a subpopulation of myelinated mechanoreceptors^11^. While data suggest c-Kit has roles in somatosensory neuron survival/differentiation or in nociceptive sensitization, its specific role in somatosensory cell biology and function is still largely unsettled^13-15^. The c-Kit ligand, SCF, is expressed in the skin of mammalian adults, suggesting a potential role in maintenance or promotion of cutaneous sensory axon density. Similar to the c-Kit receptor, zebrafish have two paralogues of SCF: *kit ligand a and b* (*kitlga/b)*, each of which primarily activates and binds its respective receptor due to biochemical selectivity^43^. At larval stages in the trunk, *kitlga* is expressed in the developing sense organs and pigment precursors, whereas *kitlgb* is expressed in the skin, similar to mammalian SCF expression^41^. To explore the effect of skin-produced Kitlgb on cutaneous axon density, we cloned *kitlgb* and generated a plasmid expressing *kitlgb* with a cleavable membrane-localized mCherry under the *keratin-4* (*krt4*) skin promoter (*krt4:kitlgb-p2a-mCherryCAAX*). Microinjection of this plasmid into *RB:GFP* embryos at one-cell stage produced larvae with mosaic patches of mCherry+ skin cells (Fig. 5A,B). Misexpression of Kitlgb in these regions induced dramatic, significant local increases in RB axon density compared to directly adjacent, mCherry-regions (Fig. 5C, Kitlgb-regions = 20.4±3.5 % axonal area, Kitlgb+ = 40.9±3.5). Together with receptor loss-of-function data, this suggests cutaneous expression of Kitlgb promotes increased somatosensory axon density in a highly localized manner.

**Figure 5:**
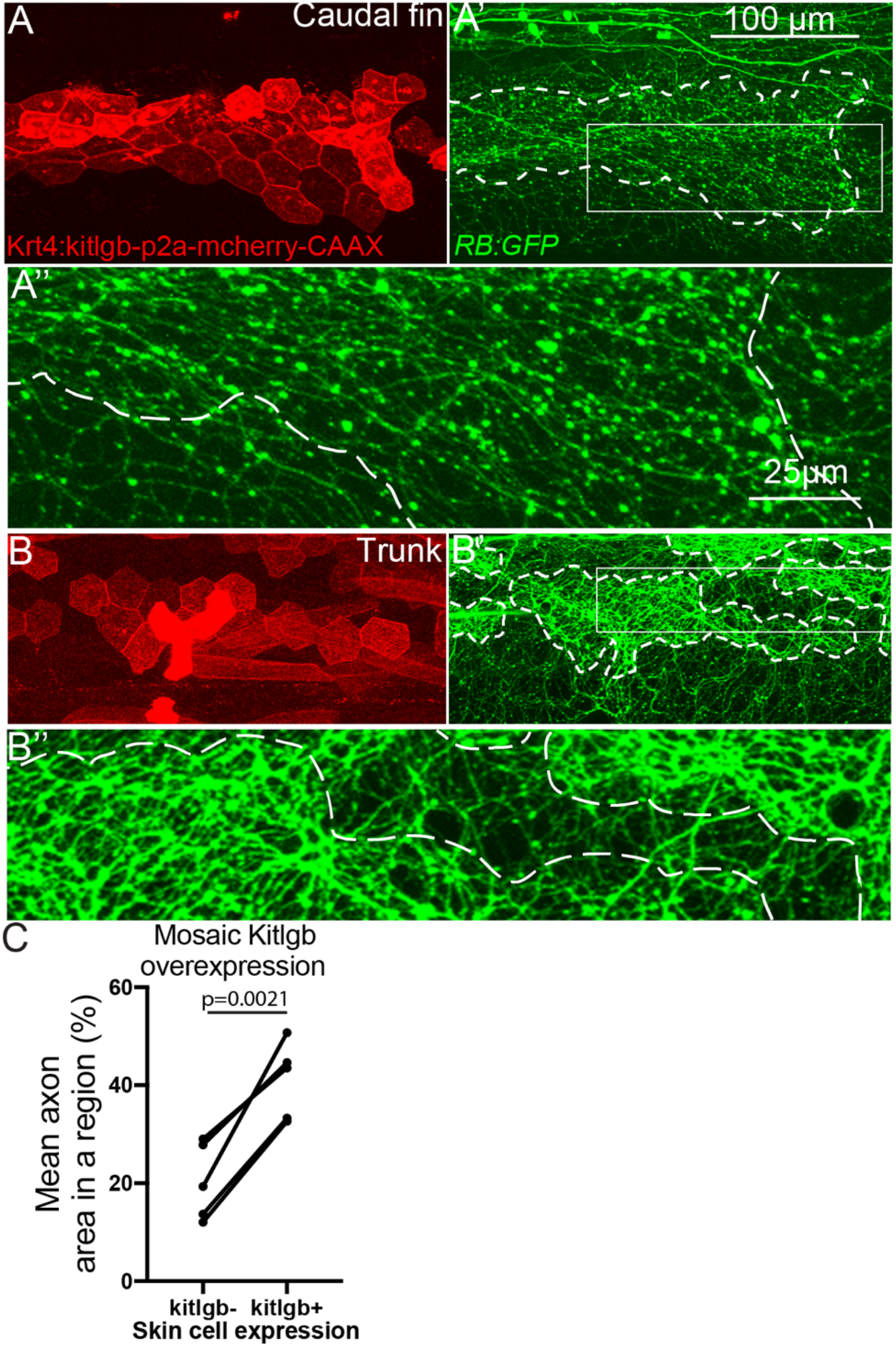
Overexpression of Kit ligand B increases local axon density. (A,B) Live imaging at 4 dpf of larvae injected with plasmid expressing *kitlgb* under a skin promoter. (A’,B’) RB axons, labeled by *RB:GFP*, are significantly denser in regions directly under dermal cells overexpressing *kitlgb* (white dashes indicate Kitlgb/mCherry+ regions). (A”,B”) 4x magnified insets of A’ and B’. (C) Quantification of axon density within individual larvae mosaically expressing Kitlgb, analyzed by paired T-test. Comparison of average axonal area of adjacent regions either mCherry positive or negative within individual larvae found significant increases in RB axon density in mCherry+ regions.

### Src is downstream of Kitb and mediates cutaneous axon density

There are several major downstream signaling pathways of activated Kit receptor, including phospholipase C (PLC) enzymes, phosphoinositide-3 kinase (PI3K), and Src^17^. To test if Src/SFK activity is a regulator of axonal density in this context, we treated *RB:GFP* larvae with saracatinib (30 μM), a potent inhibitor of SFKs, from 72-120 hpf. Saracatinib treatment induced significant loss of RB axon density similar to the other MKIs (Fig. 6B). Notably, treatment of *kitb* mutants with saracatinib did not exacerbate the loss of axon density (Fig. 6C), suggesting Src/SFKs are downstream of Kit signaling in this context. By contrast, the PLC inhibitor U73122 or PI3K inhibitor LY294002 from 72 to 120 hpf did not affect axon density (Suppl. Fig. 3).

**Figure 6:**
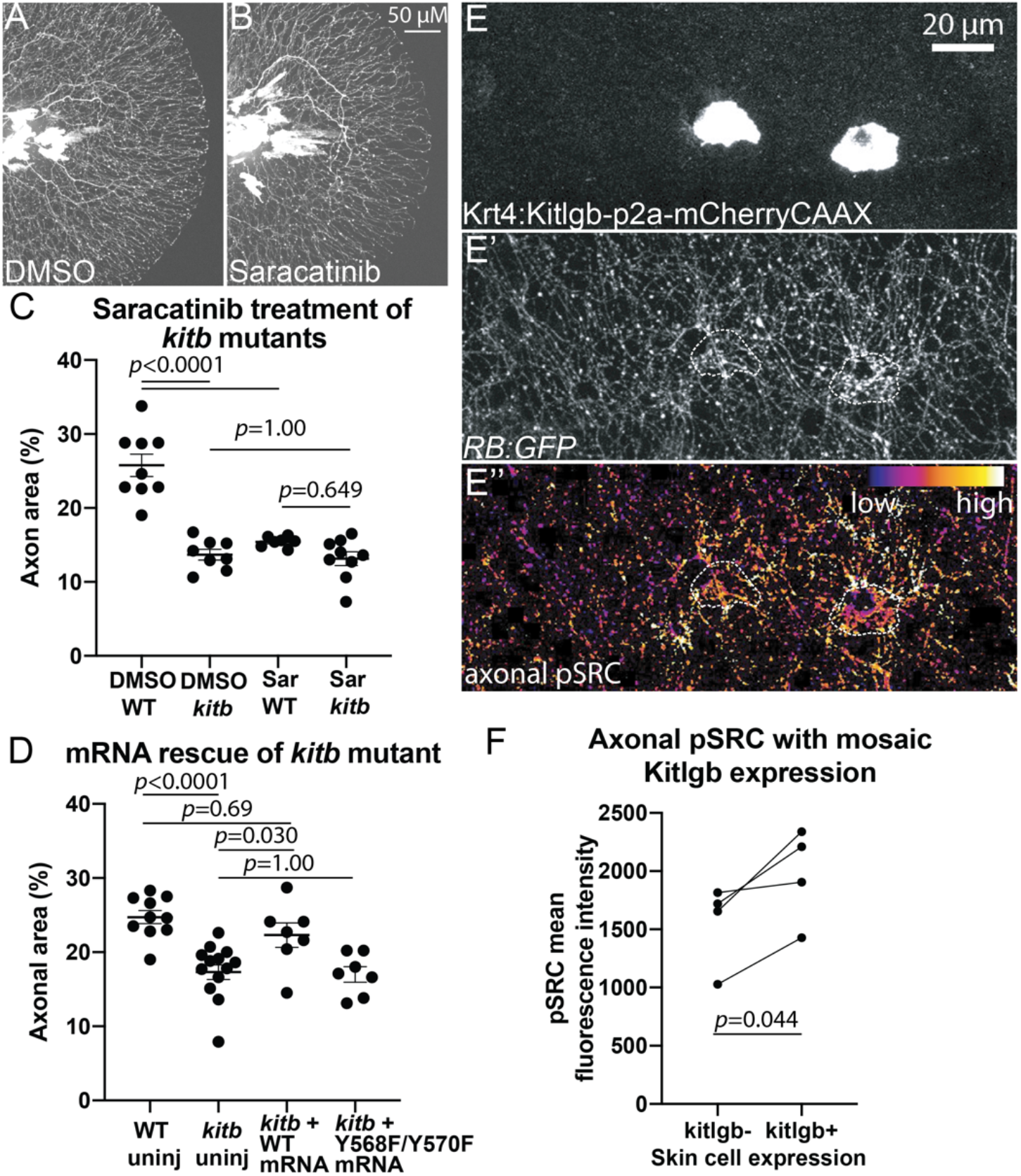
Src regulates axon density downstream of Kitb. (A,B) Lateral images of live 5 dpf *RB:GFP* tails after 48 h treatment with DMSO or saracatinib (30 μM). (C) Quantification of saracatinib treatment of *kitb* mutants, analyzed by two-way ANOVA with post hoc Tukey’s HSD test. DMSO wildtype=25.8±1.5 %area, DMSO *kitb*=13.7±0.7, saracatinib wildtype=17.2±1.2, saracatinib *kitb*=13.2±0.9. (D) Quantification of axon density of *kitb* mutants injected with wild-type or *kitb*^*Y568F,Y570F*^ mRNA (“Src-”), analyzed by two-way ANOVA with post hoc Tukey’s HSD test. WT uninjected=24.7±0.9 %area, *kitb* mutant uninjected=17.3±1, *kitb* mutant + WT *kitb* mRNA=22.3±1.6, *kitb* mutant + Y568F/Y570F *kitb* mRNA=17.0±1.1. (E-E’’) Example of Krt4:Kitlgb-p2a-mCherryCAAX-injected 3 dpf larvae immunostained for pSFK. Axonal density and pSFK staining are increased in regions near Kitlgb expression (white dashes). (F) Quantification of mean axonal pSFK intensity in Kitlgb+ vs. Kitlgb-regions within 3 dpf larvae, analyzed by paired T-test. Error bars represent SEM.

Src directly binds activated Kit receptor at two specific tyrosine phosphosites in mammals, Y567 and Y569^17^, which in turn activates Src. To test if these phosphosites were required for Kitb’s role in regulating RB axon density, we generated a Kitb receptor with phospho-dead mutations at both the zebrafish corresponding phosphosites (Y568F/Y570F) and attempted to suppress the axonal phenotype in *kitb* mutants. Unlike WT *kitb* mRNA, injection of *kit*^*Y568F,Y570F*^ mRNA did not have any significant effects of axon density in *kitb* mutants at 3 dpf (Fig. 6D). These data show that Src is a downstream target of Kit signaling that is critical for its role in regulating cutaneous sensory axon density.

We next asked if Kitlgb activation of Kitb induced changes in local axonal pSFK levels. Kitlgb was mosaically misexpressed in *RB:GFP* as described above and embryos were fixed at 3 dpf and immunostained for pSFK (Y418, Invitrogen). As previously reported, pSFK staining was highly enriched in skin cells and skin cell junctions^44^. To specifically identify pSFK activity in somatosensory axons, we created a 3D axonal volume in Imaris (Bitplane) using the GFP+ axons of the *RB:GFP* transgenic and isolated solely the pSFK signal within axons. RB axons in the RFP+, Kitlgb-expressing regions had significantly higher pSFK staining compared to adjacent RFP-regions, indicating local Kitb signaling increases SFK activity within axons (Fig. 6E,F, *kitlgb-p2a-mCherry-CAAX* positive=1970±202 mean fluorescence intensity, negative=1555±179). This indicates Kitb signaling results in local increases in both axon density as well pSFK activity.

### MKIs reduce SFK activity in distal RB axons

Because SFK activity functions downstream of Kitb in axons, we asked if MKI-mediated inhibition disrupts pSFK activity in distal axons. Acute treatment with saracatinib (7 h, 30 μM) reduced pSFK staining throughout the tail, including the axons (Suppl. Fig. 4). Acute treatment for 7 h with sorafenib (7.5 μM), which does not directly inhibit Src, did not strongly affect the skin pSFK staining but induced significant reduction in distal axonal pSFK staining (Fig. 7B”). As expected, similar treatment with ponatinib (5 μM) or dasatinib (2 μM), both of which target Src in addition to Kit, led to reduction of pSFK staining throughout the tail tissue (Fig. 7C’,D’), including significant loss of pSFK staining specifically in distal RB axons (Fig. 7C”,D”). These results indicate there is normally SFK activity in the distal axons at this time period and treatment with any of these MKIs can significantly reduce this activity. Taken together, our results suggest inhibition of Kitb reduces axonal Src activity that, in turn, disrupts normal cutaneous axon density and maintenance.

**Figure 7:**
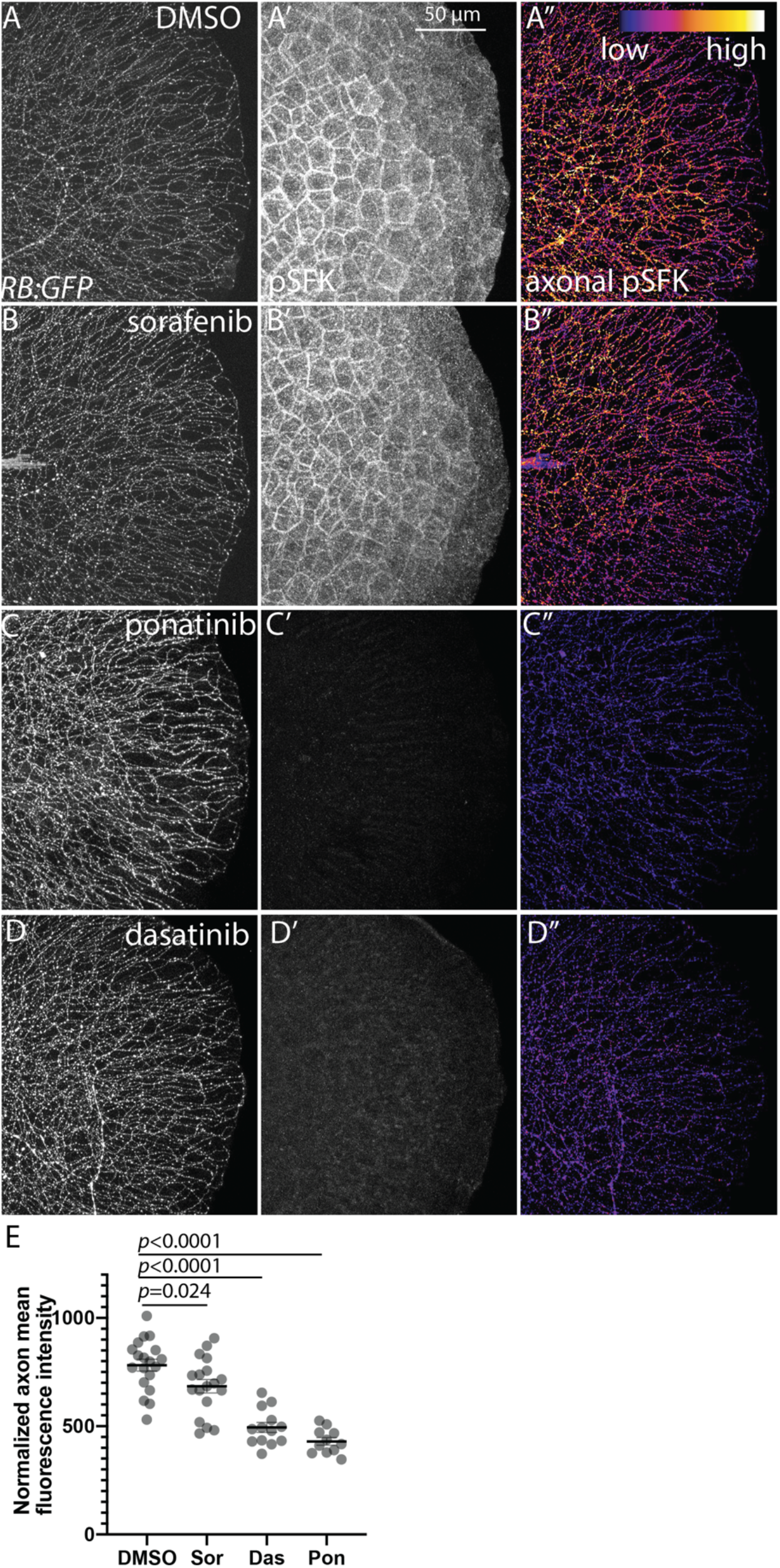
MKI treatment reduces axonal SFK activity. (A-D) Lateral view of fixed tails from *RB:GFP* larvae treated with DMSO, sorafenib, ponatinib, or dasatinib for 7 h at 3 dpf. (A’-D’) Total pSFK immunostaining. (A’’-D’’) Axonal-only pSFK staining. Sorafenib reduces axonal pSFK staining but does not notably affect other pSFK staining. Ponatinib and dasatinib abolish nearly all pSFK staining throughout the tail tissue. (E) Quantification of axonal pSFK staining following acute drug treatment, analyzed by one-way ANOVA with post hoc Dunnett’s test vs. DMSO controls. DMSO= 782±27 A.U., sorafenib=685±31, ponatinib=495±23, dasatinib=433±17. Error bars represent SEM.

## DISCUSSION

Here, we begin to address the cellular and molecular causes of MKI-induced peripheral neurotoxicity that are currently not understood. We observe *in vivo* that treatment of larval zebrafish with several neuropathic MKIs induces axon terminal die-back and loss of axon density, similar to other distal-to-proximal sensory DIPNs. We also find that Kit and Src are major drug targets underlying this neurotoxicity. Additionally, loss of Kit signaling in *kitb* mutants exacerbates vincristine neurotoxicity, suggesting Kit inhibition may enhance DIPNs in combination with non-MKI treatments. The peripheral neuropathic effects of these MKIs were also observed in SCF-responsive mouse DRG neurons, suggesting this Kit-mediated neurotoxicity is conserved in mammals. Finally, we demonstrate a novel role for Kit signaling in regulating cutaneous sensory axon density in a highly localized manner, with Src as a major mediator of this role. By characterizing several neurotoxic MKIs and identifying their molecular targets, we established larval zebrafish as a powerful tool to study MKI DIPNs and explore novel neurobiological roles of signaling pathways in somatosensory neuron development and disease. Our data provide new insight into the cellular mechanisms of MKI-induced neuropathy and lay the groundwork for studying other MKI DIPNs and the novel neurobiological roles of Kit-SFK in axon development and health.

### MKIs induce axonal die-back

To characterize the cellular mechanisms of MKI-induced sensory neurotoxicity, we observed distal somatosensory axons *in vivo* both during and after drug treatment. Vincristine is a well-established, neurotoxic chemotherapy drug that induces distal sensory axon die-back, resulting in reduced cutaneous axon density. Similarly, we observe treatment with three MKIs induced significant reduction in cutaneous axon density of the tail. Though the larval caudal tail skin is fully-innervated with dense axon arbors at the time of drug treatment (72 hpf), it is possible that reduced axon density after drug treatment is due to defects in axon growth or arborization in the developing larva. However, in wildtype larvae we find no significant changes in distal axon density during the treatment/imaging window of 3 to 5 dpf, suggesting drug-induced changes in axon density are not due to growth inhibition. Furthermore, time-lapse imaging at the onset of drug treatment demonstrates individual axon terminals significantly retracting from their initial starting location. Based on the striking similarity of the vincristine and MKI-induced phenotypes and the observed axonal retraction and loss of density, we conclude distal axonal die-back is a major cause of sorafenib, ponatinib, and dasatinib sensory neurotoxicity.

### Kit signaling mediates cutaneous innervation

c-Kit receptor is expressed in a subset of mammalian embryonic and adult somatosensory neurons, predominantly in unmyelinated, nociceptive c-fibers^10,11,13^. Expression of its ligand, SCF, is detected in mouse embryonic and adult skin as well as larval zebrafish skin^41^. Mouse mutant studies found loss of Kit signaling may affect the identity/number of some DRG neurons, hamper DRG axon regeneration, and alter responses to certain stimuli^11,13,15^. However, the exact role of Kit signaling and the relationship between the ligand/receptor expression pattern in somatosensory neuron development and function is still not fully elucidated. We find that *kitb* homozygous zebrafish mutants do not have significant loss of somatosensory neurons during development; however, they do display significant reductions in distal cutaneous axon density. Treatment of larvae well after the skin is fully innervated (3 dpf) with Kit inhibitors induces a similar reduction in axon density, suggesting this is not simply a developmental phenotype. The RB neurons affected in both cases are polymodal, unmyelinated somatosensory neurons, similar to the population of DRG neurons that express c-Kit in mammalian models and are affected in *Kit* mutant mice. We also observe reductions in neurite density of SCF-grown mammalian DRG cultures treated with MKIs, suggesting that this neuronal response to Kit inhibition is conserved in mammalian somatosensory neurons. We find that local misexpression of Kitlgb induces a dramatic, highly localized increase in cutaneous axon density. Taken together, our data suggest a critical role for Kit signaling in modulating cutaneous axon innervation and density.

The expression of Kit receptor in somatosensory neurons and the consistent expression of Kit ligand throughout the adult mouse and larval zebrafish skin suggests a constant role for Kit signaling in promoting and/or maintaining cutaneous axon innervation. We propose that the constant presence of Kit ligand induces proper axonal arborization and density levels in the skin. We observe in zebrafish that somatosensory terminals are highly dynamic, and Kit signaling may promote axon terminal outgrowth that counteracts potential retraction or loss of arborization. It may also be that SCF/Kit signaling promotes regeneration of fine somatosensory axon terminals. The cutaneous axon arbors of small-fiber, unmyelinated axons are intricate and delicate and may be prone to damage from even minor mechanical or other insult. In this case, these terminals would require consistent maintenance and regrowth and the presence of SCF throughout the skin may drive this via through Kit signaling. In either case, we provide *in vivo* evidence that Kit signaling is an important regulator of cutaneous axon density and its specific effects on sensory axon behavior bear further study. Furthermore, Kit receptor or its downstream pathways may be a candidate for future therapies attempting to ameliorate distal axonopathies in DIPNs and other diseases.

### Kit signaling in sensation

Changes in Kit signaling can affect responses to both noxious and non-noxious stimuli in mammalian models. *Kit* null mutant mice demonstrated a reduced response to noxious heat stimuli (thermal hypoalgesia) and an increased response to punctate mechanical stimuli^11^. Treatment with a Kit inhibitor recapitulated the thermal hypoalgesia phenotype but not changes in mechanical response, whereas intraperitoneal injection of SCF induced thermal hyperalgesia. Mice homozygous for the *Kit*^*W-v*^ hypomorphic allele displayed no significant change in thermal or mechanical responses; however, they did have a reduced long-term response to noxious subcutaneous formalin injection^13^. Intrathecal injection of SCF induced allodynia and an enhanced response to radiant heat stimuli but did not significantly change response to noxious heat stimuli^14^. Why genetic loss of Kit may affect responses to both noxious mechanical and thermal stimuli whereas pharmacological inhibition/activation of Kit signaling affects just responses to noxious thermal stimuli is unclear. Regardless, modulating Kit signaling affects the responses of polymodal, unmyelinated somatosensory neurons, even if the mechanisms by which Kit signaling affects responses to these stimuli is largely unknown.

Somewhat in contrast to these results, we observe loss of the provoked escape response to mild mechanical stimuli with treatment with both sorafenib and vincristine. It is possible these changes in mechanical stimulus response in zebrafish are due to the structural damage/changes to cutaneous axon density. It is unknown in other studies if the transient treatment of adult mice with SCF or Kit inhibitor was sufficient to induce similar axonal changes. It may be that Kit signaling has early roles in affecting differentiation or acquisition of specific modalities in subpopulations of somatosensory neurons and later has a role in maintaining functionality or modulating signal transduction relevant to specific somatosensory modalities.

### Kit receptor mediates MKI neurotoxicity

The incidence and severity of MKI-induced peripheral neuropathies varies from compound to compound, though the cause of this is unclear. Likely, changes in neurotoxic potential are due to the specific inhibition profile of each drug. To date, the specific kinase targets of MKIs that may induce neurotoxicity are unknown. We found that transient knockout of *kitb* reduced distal somatosensory axon density similar to sorafenib, ponatinib, and dasatinib treatments, but knockout of Ret, another RTK target expressed in RBs, had no effect. We observed that application of all these MKIs on *kitb* mutants did not enhance the axon density loss phenotype. Notably, vincristine, which does not target a specific kinase/RTK, did exacerbate the loss of axon density observed in *kitb* mutants. This suggests that loss of Kit signaling may potentially aggravate neurotoxicity of other compounds or disease states in somatosensory neurons. Further examination of the effect of Kit inhibition in contributing to neurotoxicity in combination drug therapies could provide information highly relevant to drug development and clinical decision-making. Based on its role in promoting cutaneous axon innervation, activation of the Kit signaling pathway may also provide a potent therapeutic avenue in treating other DIPNs or distal axonopathies.

### Src activity downstream of Kit signaling

Src is one of many downstream effectors of activated Kit receptor. Src/SFKs regulate axon outgrowth and pathfinding in several vertebrate systems, including *Xenopus*, chick, and mouse spinal neurons^18,20,45,46^ and chick retinal neurons^19^. SFKs also promote axon regeneration in peripheral sensory neurons in zebrafish DRGs, rat sciatic nerve, and *C. elegans* motor neurons^21-23^. In these contexts, Src/SFKs affect axon growth and steering by regulating formation and localization of filopodia and lamellipodia within growth cones, affecting motility and direction^18,47^. Though Src/SFKs are an established target of activated Kit receptor, the two have not been linked in the context of axon biology. Our results demonstrate axonal SFK activity is reduced by pharmacological inhibition of Kitb signaling and promoted locally through activation of Kitb. Additionally, Src-binding phosphosites on Kitb are required for its role in regulating cutaneous somatosensory axon density. These results suggest that Kitb/Src interaction is critical for its role in somatosensory axons and that modulation of Kitb signaling affects local axonal levels of pSFK. Based on the dynamic nature of these distal axon terminals, we suggest that activation of SFKs through Kitb maintains axon extension and aids in pathfinding as axons continue to branch and elaborate throughout the skin.

### Src as a mediator of MKI neurotoxicity

Because SFK functions downstream of Kitb signaling in axons, we hypothesized inhibition of SFKs should affect axons similarly to the other MKIs. Treatment with saracatinib induced loss of axonal density similar to other MKIs but did not exacerbate the *kitb* axon loss phenotype. This suggests SFKs are the primary downstream target of Kitb mediating its neurotoxicity and the neurotoxicity of ponatinib and dasatinib may due to direct inhibition of both Kit signaling and Src activity. These data highlight the importance of the Kit-SFK signaling axis in this MKI neurotoxicity and provide a variety of molecular therapeutic targets for future study in peripheral neuropathy and axon regeneration.

Interestingly, inhibition of SFKs can ameliorate long-term neuropathic pain in rodent models after formalin injection or peripheral nerve constriction. *N*-methyl-D-aspartate receptor (NMDAR) activity in spinal neurons is a contributor to neuropathic pain and SFK activity in these neurons enhances NMDAR activity^48^. Long-term activation of SFKs in spinal microglia may also contribute to neuropathic pain from chronic formalin injection^49^. In these cases, continued activation of SFKs is associated with continued neuropathic pain. However, SFK activity in growth cones and glia is also critical for peripheral axon regeneration^22,23^. Based on our observations, it is likely that inhibition of SFK activity in our model is inducing neurotoxicity and changes in axonal architecture by impeding its role as a promoter of axon outgrowth and regeneration. It is possible that reduction of SFK activity in the short-term contributes to distal axon die-back but that long-term reduction of SFK activity has a role in suppressing neuropathic pain after the initial insult and/or regeneration is finished. SFKs, particularly Src, have functions in almost every cell type. Thus, wholesale alteration of systemic SFK activity can have a variety of disruptive adverse effects. In contrast, modulating SFK activity through Kit signaling may offer a therapeutic opportunity to target SFKs in a cell-type specific manner to ameliorate the initial neurotoxic insult and also address SFK-mediated pain.

Characterizing the cellular pathophysiology of MKI neurotoxicity and identifying the tyrosine kinase targets that underlie it are critical to addressing this clinical issue. As the number of MKI drugs and their approved clinical applications grow, so does the prevalence of MKI DIPNs and their detrimental effect on patients. Understanding the neurobiological bases of MKI-induced neurotoxicity can inform future MKI drug design and clinical decision making and also provide therapeutic points of intervention to ameliorate both MKI-induced or other peripheral neuropathies. Here we establish larval zebrafish as a strong *in vivo* model for exploring the neurotoxic effect of MKIs and identifying the underlying molecular targets. We provide evidence that several MKIs induce distal axonal die-back and this effect is mediated by Kit signaling and SFK activity. These findings set the stage to identify and characterize other neurotoxic MKIs, elucidate the specific cellular and molecular mechanisms of Kit-SFK signaling in axon development and behavior, and explore how these signaling pathways may interact with other peripheral neuropathic compounds and disease states.

## METHODS

### Zebrafish husbandry

Adult zebrafish were maintained at 28.5°C as previously described^50^. Embryos were derived from natural matings or *in vitro* fertilization, raised in embryo media, and developmentally staged^51^. Strains utilized were ^*^AB, *SAIGFF213A*^*30*^;*UAS:GFP* (“*RB:GFP”)*, and *kitb*^*sa1534*^ mutant (obtained from Zebrafish International Resource Center, Eugene, OR, USA).

### Genotyping protocols

*kitb*^*sa1534*^ mutants were genotyped by extracting genomic DNA from tail clips or whole larvae, then PCR amplifying with *Kitb F dCAP* and *Kitb R dCAP* primers that introduce a Hind III restriction site only in the amplicon of a mutant allele. PCR products were digested with Hind III-HF (New England Biolabs) for 4 h at 37º C, and run out on a 2% agarose TBE gel to resolve digested bands. Uncut amplicon is 399 bp, whereas Hind III digests mutant amplicon in 370 and 29 bp fragments.

### CRISPR-Cas9-mediated genetic knockdown

sgRNAs for CRISPR-Cas9 knockdown of *kitb, ret, pdgfra*, and *pdgfrb* were designed (Table 2) and injections were performed as previously described^52^ into *RB:GFP* embryos. CRISPR efficiency was evaluated by isolating genomic DNA from 3 dpf larvae, amplifying regions surrounding CRISPR target sites with respective CRISPR F and R primers (Table 2) and incubating amplicons with restriction enzymes that have cut sites adjacent to CRISPR targets (*kitb* sgRNA: BslI; *ret*: BslI; *pdgfra*: DraI; and *pdgfrb*: BstNI). Digested amplicons were visualized by gel electrophoresis to ensure at least 75% of CRISPR-Cas9-injected larvae displayed total or significant loss in amplicon digestion, indicating efficient CRISPR-Cas9-mediated knockdown. At 3 and 5 dpf, *RB:GFP*+ larvae that had obvious necrosis or body morphology defects were removed and tail tips of injected and uninjected larvae were rapidly imaged using the VAST-spinning disk microscope imaging system.

### Drug treatment

Sorafenib, ponatinib, dasatinib, saracatinib, U73122, and LY294002 were obtained (Selleck Chemicals, Houston, TX, USA) and dissolved in DMSO to stock solutions of 10 mM and stored at -80º C. For larval treatment, drugs were diluted and mixed in embryo medium containing 1% DMSO and used to replace larval media. Unless otherwise noted, larvae were treated with drugs from 72-120 hpf and larval media was replaced at 96 hpf with fresh drug/vehicle-containing media.

### Plasmids, labeled constructs

To obtain template for cDNA cloning, total RNA was isolated from wildtype ^*^AB embryos using Trizol reagent (ThermoFisher Scientific). First-strand cDNA synthesis was performed on 1 µg of total RNA using oligo dT primers and Superscript III reverse transcriptase (Invitrogen). Full-length zebrafish *kitb* and *kitlgb* were amplified with *kitb* and *kitlgb clone p221* F and R primers, respectively, and recombined into the Gateway middle entry vector pDONR221 using previously described methods^53^ to generate *pME*:*kitb* and *pME:kitlgb* plasmids. The double phosphosite mutant construct *pME:kitb*^*Y568F/Y570F*^ was made by PCR using *pME:kitb* as a template and the *kitb 568/570Y-F IVA* F and R primers with the Phusion High-Fidelity PCR Kit (ThermoFisher Scientific) followed by *in vitro* assembly in Top10 competent cells (ThermoFisher Scientific) as previously described^54^.

Final plasmids were generated using the Gateway cloning method with pDestTol2pA2 destination vector and Tol2Kit plasmids unless otherwise noted^53^. The *krt4:kitlgb-p2a-mCherryCAAX* plasmid was made by recombining *p5E:Krt4*^*55*^, *pME:kitlgb*, and *p3E:p2a-mcherryCAAX-polyA. CMV/Sp6:kitb-p2a-mcherry* and *CMV/Sp6:kitb*^*Y568F/Y570F*^*-p2a-mcherry* plasmids for mRNA synthesis were made by recombining *p5E:CMV/Sp6, pME:Kitb* or *pME: kitb*^*Y568F/Y570F*^, and *p3E:p2a-mcherry-polyA. UAS:mCherry* was made by recombining *p5E:UAS, pME:mCherry*, and *p3E*:*polyA. krt4:kitlgb-p2a-mCherryCAAX* plasmid was microinjected at 20 pg/embryo and *UAS:mCherry* plasmid was microinjected at 10 pg/embryo. *Kitb-p2a-mcherry* and *kitb*^*Y568F/Y570F*^*-p2a-mcherry* mRNAs were synthesized from *CMV/Sp6:kitb-p2a-mcherry* or *CMV/Sp6:kitb*^*Y568F/Y570F*^ using SP6 mMessage Machine (Life Technologies) and microinjected at 25 pg/embryo.

### Primers and oligonucleotides

**Table 2:**
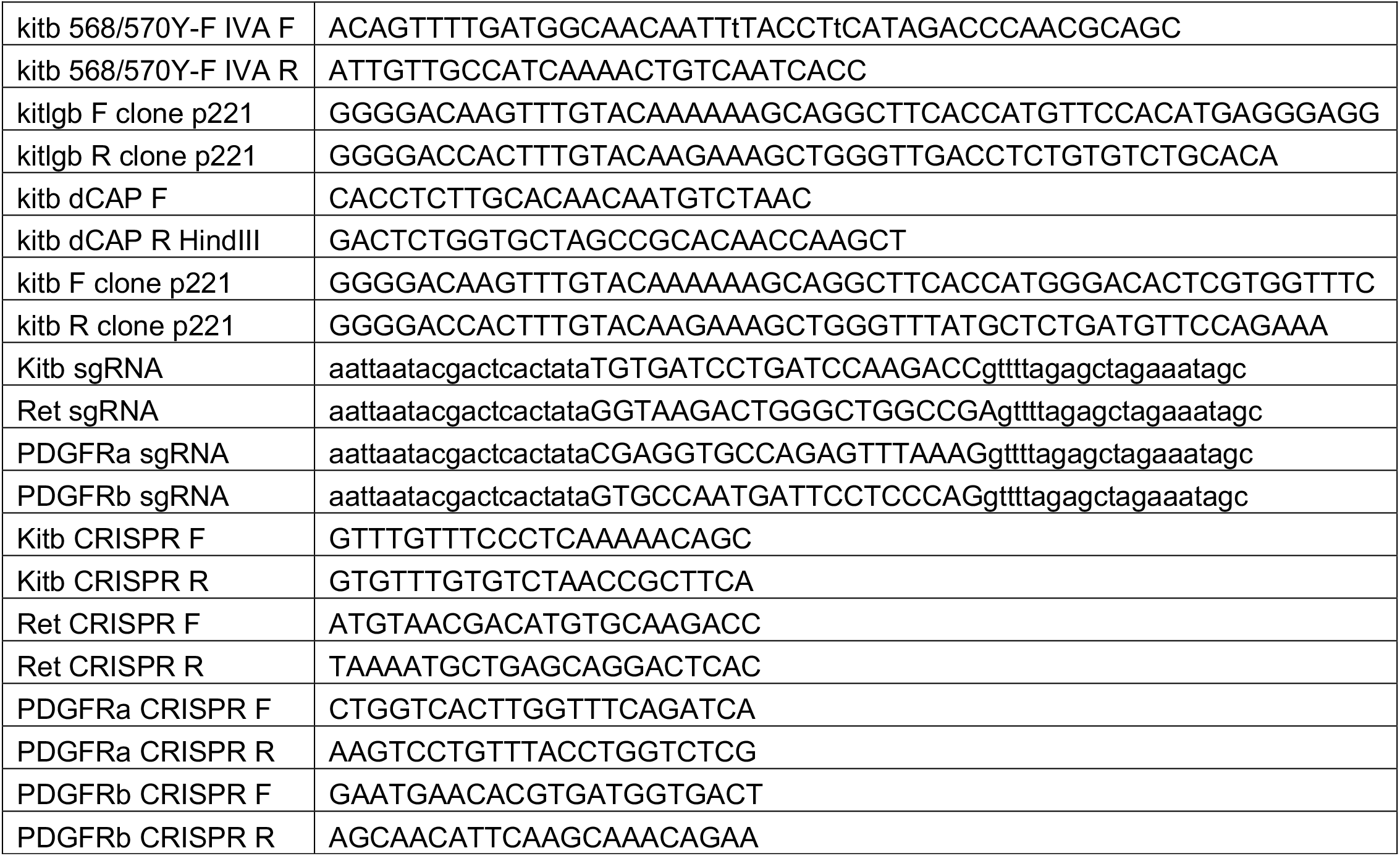
Primers and oligonucleotides

### Zebrafish immunostaining

Larvae were anesthetized in 0.02% tricaine (MS-222; Sigma) in embryo medium, fixed in 4% formalin in 1x PBS for 1 h at RT and then 4º C overnight with one exception: larvae stained for α-pSFK Tyr418 were fixed in 1.5% formaldehyde in 0.1 M PIPES, 1.0 mM MgSO_4_ and 2 mM EGTA for 1 h at RT, then 4º C overnight. Larvae were rinsed 3x with 1x PBS and permeabilized by washing in distilled water for 5 min, followed by 3x washes in 1x PBS + 0.1% Triton X-100 (PBTx). Larvae were incubated for 1 hour at RT in blocking solution (PBTx + 5% goat serum, 1% bovine serum albumin, 1% DMSO), then at RT in primary antibody diluted in blocking solution for 2 h followed by 4º C overnight. Primary antibodies used were α-pSFK Tyr418 (1:500, Invitrogen, 44660G, RRID:AB_1500523), α-mCherry (1:1000, ThermoFisher Scientific, M11217, RRID:AB_2536611), α-GFP (1:1500, Aves Labs, RRID:AB_10000240), α-Cleaved Caspase-3 (1:1000, Cell Signaling, 9664P, RRID:AB_2070042), and α-acetylated Tubulin (1:1000, Sigma-Aldric, T6793, RRID:AB_477585). Embryos were rinsed extensively in PBTx and incubated for 4 h at RT in Alexa488-, Alexa568-, or Alexa647-conjugated secondary antibodies diluted in blocking solution (1:1000, ThermoFisher Scientific). After rinsing in PBTx, larvae were transferred to 70% glycerol in PBS and mounted on slides with 0.13-0.17 mm borosilicate coverslips secured with vacuum grease.

### Confocal fluorescence imaging of zebrafish larvae

For live-imaging, larvae were mounted in 1.5% low melting point agarose on a glass coverslip, submerged in embryo media containing 0.02% tricaine (MS-222; Sigma) and imaged using a 40x/NA=0.8 water immersion objective on an upright Fluoview1000 confocal microscope (Olympus) with 488 and 568 nm excitation channels. To assess RB neuron apoptosis over time, RB neuron cell bodies in the dorsal spinal cord of *RB:GFP* embryos were imaged at 10 min/frame for 15 h in an ∼400 μm wide viewing window. For imaging sparsely labeled RB neurons over time and the axonal effects of Kitlgb misexpression, *RB:GFP* embryos were injected at 1-cell stage with *UAS:mCherry* or *Krt4:kitlgb-p2a-mCherryCAAX* plasmids, respectively. Larvae injected with *UAS:mCherry* that contained caudal tail axons expressing mCherry at 3 or 4 dpf were mounted, submerged in embryo media containing vehicle or drug, and imaged 5 min/frame over 10 h. *RB:GFP* larvae injected with *Krt4:kitlgb-p2a-mCherryCAAX* were screened for presence of mCherry+ skin cells and the mCherry+ regions and mCherry-adjacent regions were imaged. Immunostained larvae were imaged on the Olympus Fluoview1000 and 488, 568, and 647 nm excitation channels. For cleaved Caspase-3 immunostained larvae, a 20x/NA=0.75 air objective was employed and for pSFK immunostained larvae, a 60x/NA=1.42 oil immersion objective was used.

### High-throughput fluorescence larval imaging

For live high-throughput imaging caudal tails of wild-type, *kitb* mutant, or drug-treated *RB:GFP* larvae, we used the Large Particle (LP) Sampler and VAST BioImager (Union Biometrica) to automate the delivery of zebrafish from 96 multi-well plates to the BioImager microscope platform as previously described^56^. The BioImager platform was mounted on an upright Axio Examiner D1 (Carl Zeiss) microscope, fitted with a high-speed CSU-X1 spinning disk confocal scanner (Yokogawa, Tokyo, Japan). A 1.6X post-magnification adaptor was added to the light path combined with a C-Plan-Apochromat 10X/NA=0.5 dipping lens (Carl Zeiss) and an AxioCam 506 m CCD camera (Carl Zeiss). This setup allowed acquisition of images with X-Y dimensions of 662 µm × 492 µm, such that the majority of the capillary’s 600 µm Y-axis could be captured. The imaging system was configured with a 50 mW 405 nm laser and a high-speed emission filter wheel with 520/35 and 527/54 emission filters for the camera. For rapid image acquisition in the Z-plane, we employed a long-range, high-speed PIFOC P-725.4CD piezo objective scanner (Physik Instrumente, Karlsruhe, Germany), capable of up to 400 µm high speed travel. Z-stacks of 218 frames with a step size of 1.84 μm were acquired using Zen Blue 2.0 software (Carl Zeiss).

### Axon density quantification

Based on the large volume of data produced by high throughput, we designed a streamlined system for measuring axonal density using open-source software. We imported confocal images of *RB:GFP* caudal tails into FIJI^57^, converted images to 8-bit, and generated maximum intensity Z-projections. We then thresholded the projections, formed a rectangular 50×100 μm ROI at the distal tail tip, and measured the percentage of this ROI occupied by GFP+ axons.

To quantify changes in axonal density in cutaneous tissue misexpressing Kitlgb, we generated max-intensity Z-projections of imaged regions of *Krt4:kitlgb-p2a-mCherryCAAX*-injected larvae and made ROIs using the boundaries of mCherry+ and adjacent mCherry-regions of skin cells in FIJI. Percent area measurements of *RB:GFP-*labeled axons in adjacent mCherry+/mCherry-regions were taken as described and compared within individual larvae.

### Immunofluorescence intensity quantification

For consistency of labeling, compared larvae were processed in the same batch. To quantify axonal pSFK staining, *RB:GFP* larvae were immunostained with α-pSFK, α-GFP, and α-acetylated Tubulin (AcTu). 3D reconstructions of larvae axonal volumes were generated in Imaris (Bitplane) using GFP immunofluorescence channel signal. For quantifying pSFK changes in the caudal tail, we measured mean fluorescence intensity (MFI) of pSFK and AcTu within axons in a 50×100 μm ROI at the distal tip of the tail. To account for sample-to-sample variation in permeability and staining intensity, we normalized pSFK MFI to the ratio of the AcTu MFI of an individual larva divided by the average of the AcTu MFI within an experimental group. To prepare 2D images of pSFK immunostaining solely in axons, we generated a masked channel containing only pSFK signal within the 3D-reconstructed axons, and made max intensity Z-projections for each channel in Imaris.

To quantify axonal fluorescence intensity of pSFK staining in cutaneous regions misexpressing Kitlgb-p2a-mCherryCAAX, we identified immunostained larvae containing regions of 1-2 isolated mCherry+ skin cells. These regions (1-2 per larva) were imaged and axonal volume was reconstructed in Imaris as described. Axonal pSFK MFI was measured within mCherry+ regions and within two adjacent, mCherry-regions of comparable size. Within a single larva, MFI of mCherry+ and mCherry-regions were separately averaged and then compared.

### Axon terminal dynamics analysis

Distance of all distal axon terminals from tail edge that were within 50 μm of the distal caudal tail edge at t = 0 per labeled RB neuron were measured at t = 0 and t = 10 h and averaged for each neuron (n = 5 total neurons in ≥3 larvae per condition). For kymograph generation, analysis, and visualization, we used the MetaMorph software package (Molecular Devices, Inc.) as previously described^58,59^ with one difference: instead of analyzing movement of individual, labeled particles, we examined single axon terminals labeled by *UAS:mCherry* in *RB:GFP* larvae to visualize net movement relative to the nearest tail edge over time.

### Scoring apoptotic RB neurons

*RB:GFP+* wild-type or *kitb* mutant larvae were fixed at 30 or 48 hpf, immunostained for cleaved-Caspase 3, and imaged from otic vesicle to the end of the spinal cord. Because apoptotic and fixed neuron cell bodies did not always retain the main morphological hallmarks used to identify RB neurons, we counted in *RB:GFP*+ and cleaved-Caspase 3+ neurons in the dorsal-most portion of the spinal cord, where only RBs should be present. For time-lapse imaging of RB neuron cell bodies, the posterior-most edge of the 400 μm viewing window was placed in line with the end of the yolk extension. RB neurons were identified by spinal cord location, morphology, and axonal projections and apoptotic/dying neurons were counted. Counts per viewing window were normalized to 50 μm length for comparison.

### Plate and Coverslip Preparation for DRG culture

DRG Explants were grown on glass coverslips in 12-well tissue culture plates. The day before dissection glass coverslips were placed in wells, sterilized with 70% ethanol for 10 minutes, then the ethanol was aspirated and the coverslips were allowed to air dry. Coverslips were then washed 3 times with cell culture grade water and were incubated overnight at 4°C in 100 µg/mL poly-D lysine in cell culture grade water. Following this incubation coverslips were washed twice with cell culture grade water and were incubated at 37°C with 10 µg/mL mouse Laminin (ThermoFisher, 23017015) for 5 hours before plating. When explants were ready to plate, coverslips were washed twice with cell culture grade water and allowed to dry for 5-10 minutes before plating.

### In-vitro *DRG explant culture*

DRG explant cultures were performed on E15.5 CD1 mouse embryos. Pregnancy timing was determined by vaginal plug, counting the morning of plug presence as E0.5. Pregnant dams were euthanized according to IACUC approved methods, and the uterus was removed and placed into chilled Leibovitz’s L-15 Medium (ThermoFisher, 11415064) with Penicillin/Streptomycin (0.2%). Embryos were removed from the uterus and amniotic sacs, then were transferred to a plate containing fresh L-15/Penicillin/Streptomycin. Dorsal root ganglia (DRG) from a single embryo were isolated through fine dissection as previously described^60^. DRGs from a single embryo were transferred to a tube on ice using an FBS-coated, fire-polished glass pipette. All explants on a single plate of cultures were derived from a single embryo. Isolated DRGs were digested in 1 mL of 1 mg/mL Collagenase (Fisher, NC9693955), 1 mg/mL Dispase (Worthington Biochemical, LS02104) in L-15 media for 20 minutes at 37°C with gentle agitation every 5 minutes. 1-2 µL of DNAse (Fisher, PRM6101) was added during incubation if needed to reduce clumping of the digesting DRGs. After digestion the DRGs were briefly spun down and the supernatant was carefully removed. The DRGs were resuspended in L-15 media with 10% fetal bovine serum (FBS) and gently agitated to break up the pellet. The DRGs were briefly spun down, the supernatant removed, and resuspended in L-15 media twice to ensure removal of remaining collagenase, dispase, and DNAse. After washing DRGs were suspended in DRG plating media (Neurobasal media, ThermoFisher, 21103049), 35mM glucose, 2% B27 supplement (ThermoFisher, 17504044), 1% GlutaMAX supplement (ThermoFisher, 35050061), 1% Penicillin/Streptomycin, 50 ng/mL stem cell factor (Proteintech, HZ-1024) run through a 0.22 µm syringe filter. DRGs were triturated 5 times through a fire-polished, FBS-coated glass pipette by aspirating and dispensing the DRGs and media with the pipette gently placed against the bottom of the tube. DRG pieces were washed 5 times by allowing the tube to sit for 1 minute, allowing the large DRG pieces to settle to the bottom of the tube, then removing the supernatant and resuspending in L-15 media. After washing DRGs were resuspended in DRG plating media and were spot plated onto prepared PDL and Laminin coated glass coverslips. Spots of approximately 20 µL each containing 1-2 DRG explants were placed on each coverslip. Plates were incubated for 1 hour at 37°C with 5% CO_2_ to allow for explants to adhere to the coverslips before gently flooding the wells of the plate with DRG plating media.

After 24 hours of growth, a media change was performed with half the plated volume of media and treatment drugs were added at appropriate concentrations in DMSO. Cultures were allowed to grow for another 24 hours at 37°C in 5% CO_2_ before fixation in for 30 minutes at room temperature by adding 1 plating volume of 4% Paraformaldehyde-Krebs Solution-Sucrose fixative (4% w/v paraformaldehyde, 145 mM NaCl, 5 mM KCl, 1.2 mM CaCl_2_-2H_2_O, 1.3 mM MgCl_2_-H_2_O, 1.2 mM NaH_2_PO_4_-H_2_O, 10 mM glucose, 20 mM HEPES, 0.4 M sucrose, in H_2_O) for a final paraformaldehyde concentration of 2%. After fixation coverslips were washed 3 times for 10 minutes with PBS at room temperature without agitation before progressing to immunolabeling.

### Explant Immunolabeling

Fixed and washed explant containing coverslips were incubated for 30 minutes in blocking buffer (0.2% Triton X-100, 5% Normal Donkey Serum, 0.002% NaN_3_, 5% DMSO, in phosphate-buffered saline) at room temperature without agitation. Coverslips were incubated in blocking buffer with primary antibody (Rabbit α-β-III Tubulin, Millipore Sigma T2200-200UL, 1:1000, RRID: AB_262133) overnight at 4°C without agitation. After primary incubation coverslips were washed with PBS at room temperature for 10 minutes four times. Nuclei were labeled with Hoechst 33342 (1:5000 in PBS, Life Technologies H3570) included in the first wash after primary incubation. Coverslips were incubated in blocking buffer with fluorescent secondary antibodies (Donkey α-Rabbit 488 or Donkey α-Rabbit 546, ThermoFisher A-21206 or A-10040) for 4 hours at room temperature in the dark without agitation. Coverslips were washed three times with PBS at room temperature for 10 minutes without agitation. Coverslips were mounted to glass slides for imaging using Fluoromount-G (Fisher, OB100-01) for imaging.

### Explant Imaging and Quantification

Explants were imaged on a Zeiss AxioImagerM.2 upright microscope using a 5x objective. Quantification was performed in FIJI^57^. The outline of the dense area of cell bodies at the center of an explant was traced manually and a region of interest was generated by expanding this traced shape 50 µm and using the FIJI “Make Band” function to generate a 50 µm wide band. The image was then manually thresholded and the coverage density of the 50 µm wide band was measured.

### Statistical Analysis

Analysis was performed with Prism software (Graphpad). Specific statistical tests and post hoc tests for each data set are indicated in text and figure legends. For experiments involving two independent variables, two-way ANOVA tests were performed to test if main effects and interactions were statistically significant. If interaction was statistically significant, ANOVA was repeated with simple effects, and significance of main effects was re-evaluated and the indicated post hoc tests were performed.

## Ethics statement

All animal works were approved by and conducted according to guidelines of the Oregon Health and Science University IACUC.

