## Supplemental Figures for "c-Kit mediates cutaneous sensory axon innervation and multi-kinase inhibitor-induced neurotoxicity"

**SUPPLEMENTARY DATA**

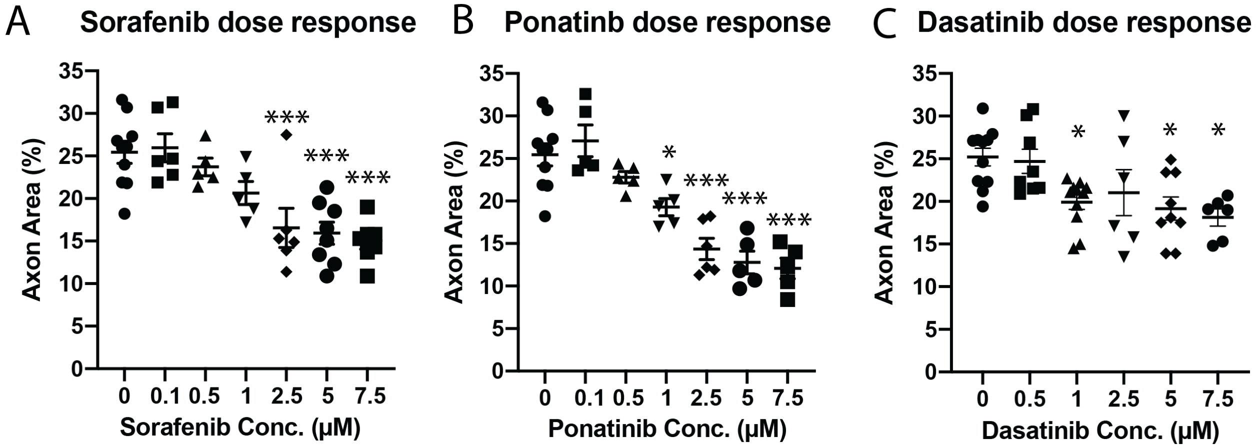


Supplementary Figure 1: Dose dependent effect on axon density with MKI treatment. (A-C) Quantification of axonal area of *RB:GFP* larvae treated at 3 dpf for 48 h with different concentrations of sorafenib (A), ponatinib (B), or dasatinib (C). All drugs show a trend of dose-dependent decrease in axonal area in response to increasing drug concentration. Error bars represent SEM, data analyzed by one-way ANOVA with post hoc Dunnett's test vs. control. *=$p<0.05$, **=$p<0.01$, ***=$p<0.001$.

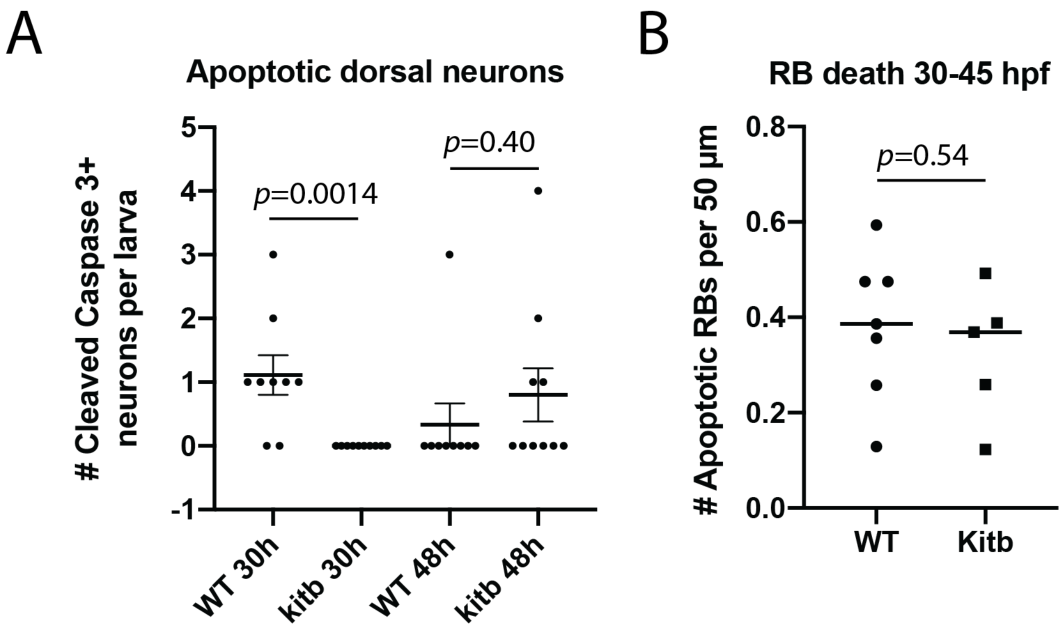


Supplementary Figure 2: Absence of RB cell death in *kitb* mutants. (A) Quantification of Cleaved-Caspase 3+ dorsal neurons in immunostained *RB:GFP* wildtype and *kitb* mutant larvae, analyzed by unpaired Student's T-test at each time point. There was no significant difference in GFP+/cleaved-Caspase-3+ dorsal neuron number in *kitb* mutants at 48 hpf compared to siblings (sibling = 0.3±0.3, *kitb* = 0.8±0.4) but at 30 hpf, there were significantly fewer of these cells detected in *kitb* mutants (siblings = 1.1±0.3, *kitb* = 0±0). (B) Quantification of time-lapse observation of RB cell death during embryonic development, analyzed with unpaired T-test. Error bars represent SEM. *RB:GFP kitb* mutants and siblings were time-lapse imaged at the mid-trunk region from 30-45 hpf. RB neurons were identified by anatomical location and morphology. There was no significant difference in the number of RB neurons undergoing apoptosis between *kitb* mutants and siblings: sibling=0.4±0.1 neurons/50µm, *kitb*=0.3±0.1.

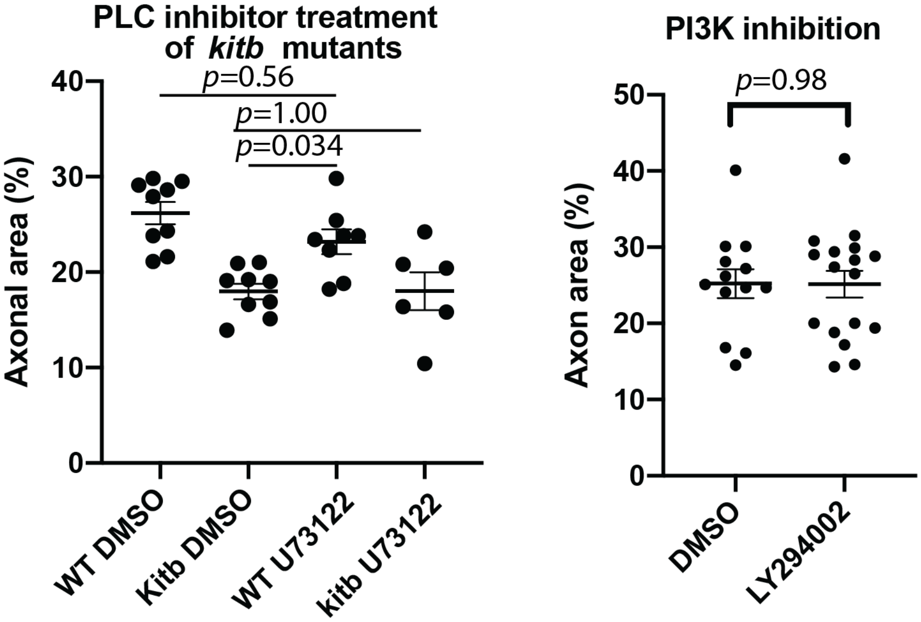


Supplementary Figure 3: Treatment of *RB:GFP* larvae with PLC and PI3K inhibitors. (A) Quantification of treatment of wild-type and *kitb* mutant larvae with 0.75 µM PLC inhibitor U73122. U73122 treatment did not significantly affect axonal area in wild-type or *kitb* mutant larvae. WT DMSO= 26.1±1.2 % area, *kitb* DMSO=18.0±0.8, WT U73122=23.1±1.3, *kitb* U73122=18±2.0. Analyzed by two-way ANOVA with post hoc Tukey's HSD test. (B) Quantification of treatment of larvae treated with 25 µM LY294002. Treatment with LY294002 had no significant effect on axonal area. Analyzed by unpaired T-test. Error bars represent SEM.

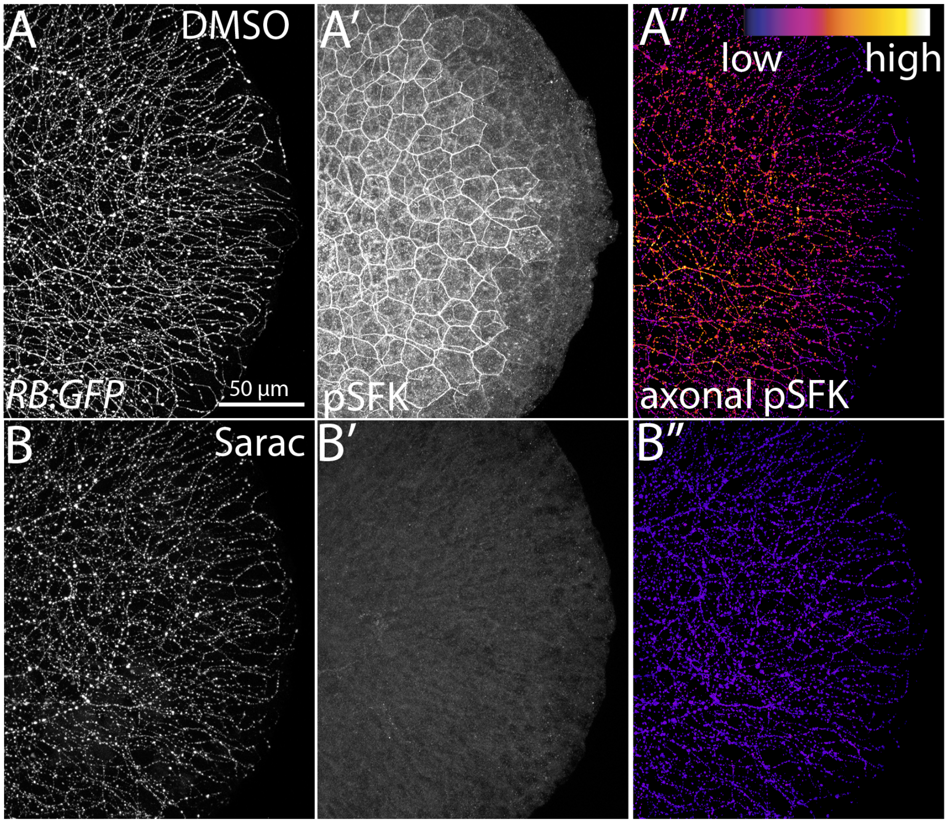


Supplementary Figure 4: Acute saracatinib treatment of *RB:GFP* larvae. (A,B) 55 hpf fixed tails after 7 h acute treatment with 30 µM saracatinib treatment immunostained for pSFK. (A) DMSO-treated larvae display strong pSFK staining in both skin and axons whereas saracatinib-treated larvae (B) lack almost any pSFK signal throughout the tail tissue.
